# Spatial modulation of dark versus bright stimulus responses in mouse visual cortex

**DOI:** 10.1101/2020.10.27.353573

**Authors:** Brice Williams, Joseph Del Rosario, Stefano Coletta, Edyta K. Bichler, Tomaso Muzzu, Anderson Speed, Lisa Meyer-Baese, Aman B. Saleem, Bilal Haider

## Abstract

A fundamental task of the visual system is to respond to luminance increments and decrements. In primary visual cortex (V1) of cats and primates, luminance decrements elicit stronger, faster, and more salient neural activity (OFF responses) than luminance increments (ON responses). However, studies of V1 in ferrets and mice show that ON responses may be stronger. These discrepancies may arise from differences in species, experimental conditions, or from measuring responses in single neurons versus populations. Here, we examined OFF versus ON responses across different regions of visual space in both single neurons and populations of mouse V1. We used high-density silicon probes and whole-cell patch-clamp recordings to assess OFF versus ON dominance in local field potential (LFP), single neuron, and membrane potential responses. Across these levels, we found that OFF responses clearly dominated in the central visual field, whereas ON responses were more evident in the periphery. These observations were clearest in LFP and subthreshold membrane potential. Our findings consolidate and resolve prior conflicting results and reveal that retinotopy may provide a common organizing principle for spatially biasing OFF versus ON processing in mammalian visual systems.

## Introduction

Visual systems respond to both increases and decreases of luminance (Schiller, 1995; Joesch et al., 2010). In mammals, stimuli that are either brighter or darker than their surroundings activate distinct retinal ganglion cells. These transmit relative luminance increases (ON responses) versus decreases (OFF responses) as action potentials to the rest of the brain (Kuffler, 1953; Field and Chichilnisky, 2007). Since retinal ganglion cells only respond to stimuli in distinct portions of the visual field, their ON versus OFF responses also implicitly carry spatial information. The spatial representation of ON versus OFF signals thus forms a critical substrate for visual perception.

Ample yet conflicting evidence indicates mammalian ON versus OFF systems are not equal. Humans detect OFF signals more quickly and more accurately than ON signals (Komban et al., 2011). Likewise, studies in macaques show that OFF signals are stronger than ON signals in primary visual cortex (V1) in layer 2/3 (Yeh et al., 2009; Xing et al., 2010). OFF responses are also stronger, faster, and more widespread in cat V1 (Jin et al., 2008; Liu and Yao, 2014). In contrast, L2/3 neuron Ca^2+^ responses in ferrets show ON dominance (Smith et al., 2015). These differences could be related to species, recording technique, or stimulus properties, particularly stimulus position in the visual field. Indeed, stimuli in the center of vision elicit OFF dominant responses found in primates and cats, whereas stimuli in the periphery elicit ON dominance in ferrets (Smith et al., 2015). Measuring electrophysiological ON versus OFF dominance in both central and peripheral visual space within species would reveal if retinotopy—a property found across all visual systems— explains these differences.

The mouse has emerged as an important tool for investigating spatial vision (Saleem et al., 2018; Wang and Krauzlis, 2018; Speed et al., 2020). The peripheral visual field of mice encompasses a large monocular region, while the central visual field spans at least 40° of binocular visual space (Wallace et al., 2013; Samonds et al., 2019). Studies of ON versus OFF signals in mouse V1 pose several unresolved questions. First, there is no agreement if mouse V1 is ON or OFF dominated. In L2/3, Ca^2+^ responses can show equal ON and OFF dominance (Smith and Hausser, 2010), strong OFF dominance (Jimenez et al., 2018), or strong ON dominance measured with voltage imaging (Polack and Contreras, 2012). Second, the laminar profile of OFF versus ON dominance beyond L2/3 remains unknown. Third, it remains unknown if ON versus OFF dominance arises from selectivity of subthreshold inputs. Combining several recording techniques in mice could help resolve these questions about the cellular, laminar, and synaptic basis of ON versus OFF dominance across the retinotopic map in V1.

Here, we measured OFF versus ON dominance in monocular and binocular mouse V1 at population, single neuron, and subthreshold levels. We first performed laminar silicon probe recordings of local field potentials and single neuron spikes in V1. We compared these results to two independent data sets that also used laminar silicon and Neuropixels probes. We then used whole-cell patch-clamp recordings to measure ON versus OFF selectivity of membrane potential. Across levels, we found strong OFF dominance in binocular V1, and weaker trends towards ON dominance in monocular V1. These findings were clearest at LFP and subthreshold levels. Our results reveal that OFF versus ON dominance in mouse V1 varies as a function of retinotopy.

## Results

### OFF dominance in binocular V1 LFP, ON dominance in monocular V1 LFP

We first measured LFP responses in awake, head-fixed mice with multi-site laminar silicon probes. White or black bars (9° wide, 0.1s duration, vertical orientation, inter-stimulus interval 0.3s) appeared one at a time on isoluminant linearized grey screens at randomly selected contrast levels and positions subtending 150° of visual space (Fig. 1A). This space spanned both the binocular and monocular visual fields. Recordings targeted to binocular V1 revealed spatially localized responses to high contrast stimuli appearing within 20° of the vertical meridian (Fig. 1B). Here, black bars elicited 48 ± 27% larger LFP responses (±SD) than white bars at these same locations in the receptive field (RF; Fig. 1D). In contrast, recordings targeted to monocular V1 revealed 31 ± 23% larger LFP responses to white rather than black bars in the RF (Fig. 1C, E).

**Fig. 1.**
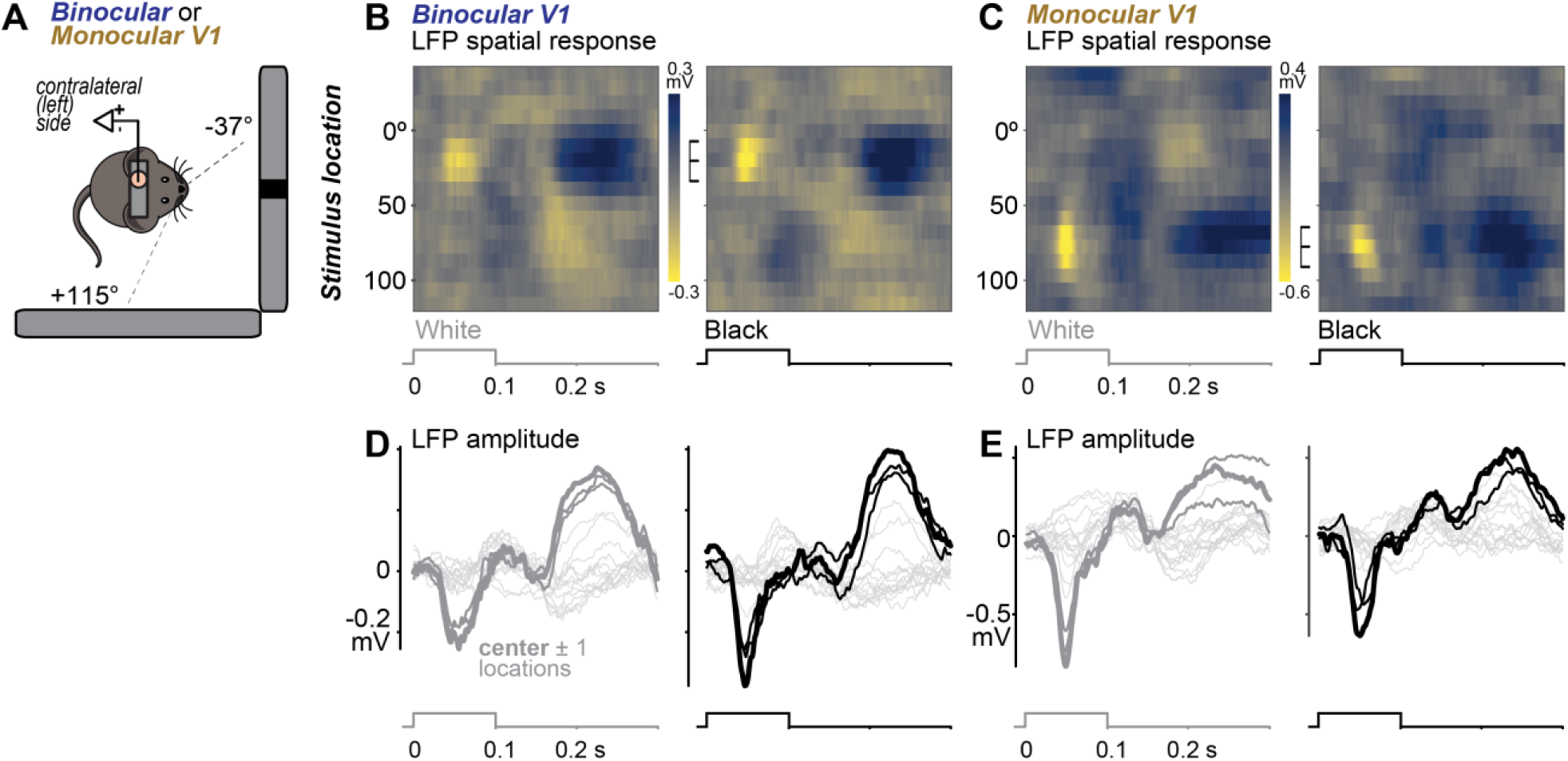
Black versus white stimulus responses in binocular versus monocular V1. **A:** Head-fixed awake mice viewed black and white bars (9° wide, full screen height) presented across 155° of azimuthal visual space. Bars appeared one at a time at randomized location, contrast, and polarity. Vertical meridian defined as 0 °, bars displayed from −37 to 115°. Neural activity recorded simultaneously with linear 32 channel silicon probe in binocular or monocular regions of primary visual cortex (V1). **B:** Example local field potential (LFP) responses in binocular V1 evoked by white (left) or black (right) bars appearing across azimuthal locations (ordinate). Median LFP responses to each bar presentation per location calculated across channels to create space-time receptive field (RF) maps. Brighter colors indicate depth negative LFP (activation). Maximum activation for black bars at 20°. Brackets span best 3 stimulus locations (center ± 1 locations) for the recording. Stimulus timing shown below maps (abscissa). **C:** Same as B, for a recording from monocular V1. Maximum activation for white bars at 77°. **D:** Median LFP responses across channels to white (left) and black (right) bars for binocular recording in B. Best stimulus location (center) defined by largest average evoked LFP response (thick trace). Responses to stimuli at the two adjacent (± 1) locations shown in thin bold traces, responses at all other locations shown in light grey. Peak binocular response 48 ± 27% (± SD) larger for black (−0.38 mV) than white (−0.26 mV) bar at same location. **E**. Same as D, for monocular recording in **C**. Peak monocular LFP response 31 ± 23% larger for white bars (−0.83 mV) than black bar responses at same location (−0.64 mV).

Binocular V1 exhibited significant OFF dominance across LFP recordings. As in prior studies, we computed the signal to noise ratio (SNR, a metric of response amplitude normalized by baseline standard deviation, see Methods) for each recording.

We quantified OFF versus ON dominance by log_10_ transforming the ratio of LFP SNR evoked by white versus black stimuli; a log ratio less than 0 indicates a response preference for black bars (OFF dominance), while a ratio greater than 0 indicates preference for white bars (ON dominance). Across all cortical layers (defined by CSD analysis, Fig. S1), LFP responses in binocular V1 showed significant OFF dominance (Fig. 2A; *p*< *0*.*001* across all layers, sign test; n = 58 recordings, 10 mice; median preferred RF locations 19.8 ± 7.1°). Opposite biases were observed in monocular V1: LFP responses across layers tended to be ON dominant, but not significantly (Fig. 2B; n = 36 recordings, 9 mice; median RF preference: 77.4 ± 8.5°). In both locations, higher contrast accentuated the luminance polarity preferences (Fig. S2), consistent with prior reports (Liu and Yao, 2014). Cumulative responses to full contrast stimuli across all layers and all recordings (Fig. 2C) showed that binocular responses were significantly OFF dominated (86% of log ratios < 0; *p* < *0*.*001*; sign test), while monocular responses were ON dominated (57% of log ratios > 0) but not significantly (*p = 0*.*38*, sign test). However, the two distributions were clearly significantly different from one another (*p* < *0*.*001;* two-sided Kolmogorov-Smirnov test).

**Fig. 2.**
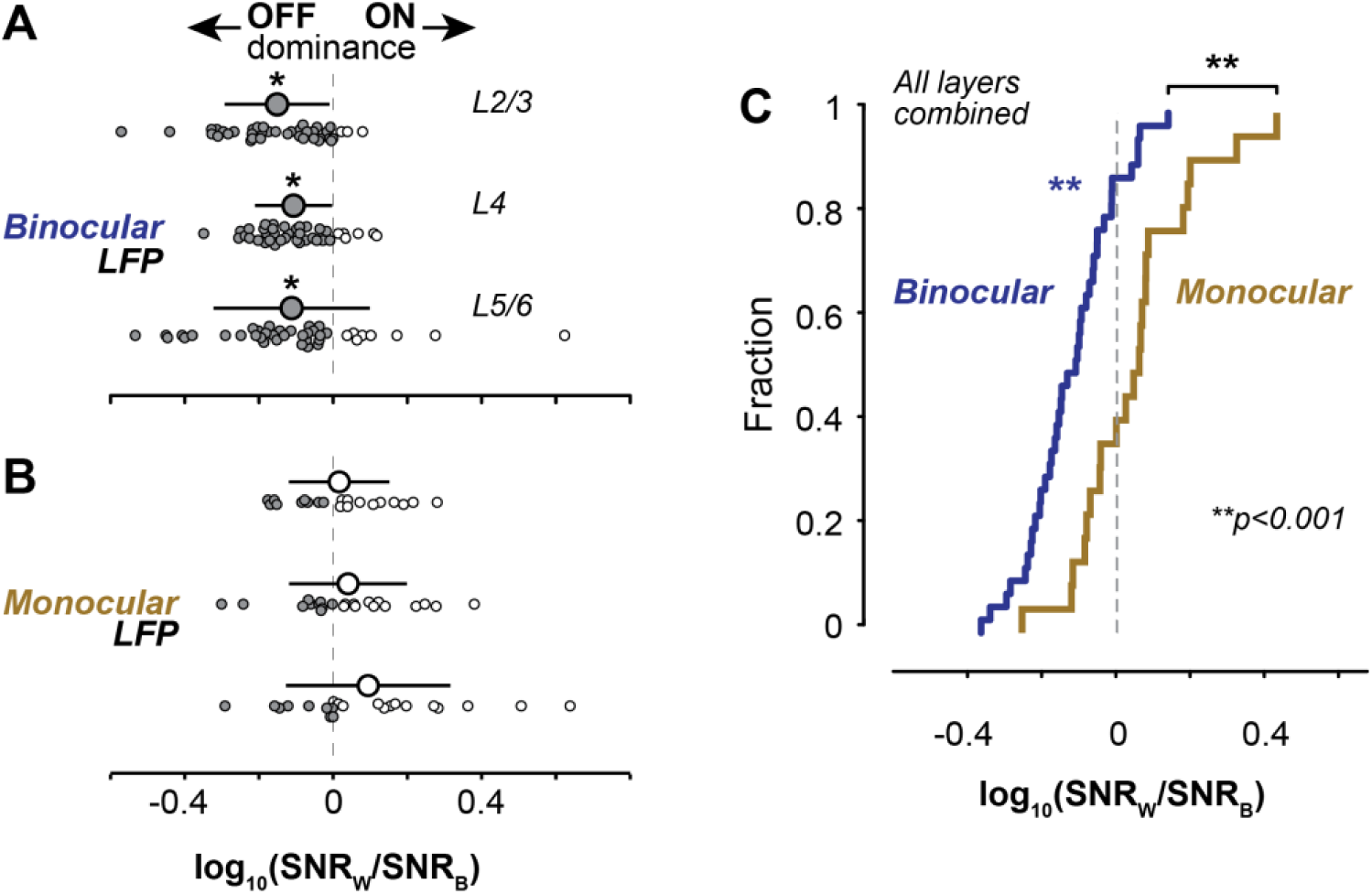
OFF dominance in binocular V1 LFP, ON dominance in monocular V1 LFP. **A:** Log_10_ ratios of LFP SNR driven by full contrast white versus black stimuli, split by cortical layer (see Fig. S1, S2). Binocular LFP responses significantly OFF dominated (log ratio < 1) across layers (L2/3: −0.15 ± 0.14, L4: −0.11 ± 0.10, L5/6: −0.11 ± 0.21; mean ± SD throughout figure; *p*<*0*.*001 for all*, two-sided sign test; n = 58 recs in 6 mice). **B**. Same as A, for monocular recordings. Monocular responses ON dominated (log ratio > 1) across layers, but not significantly (L2/3: 0.016 ± 0.14, L4: 0.04 ± 0.16, L5/6: 0.094 ± 0.22; *p = 0*.*82, 0*.*83, 0*.*52*, two-sided sign test; n = 36 recs in 7 mice). **C:** Combined across layers, binocular responses significantly OFF dominated (blue; −0.12 ± 0.12, *p*<*0*.*001*, sign test), monocular responses ON dominated (gold; 0.047 ± 0.16, *p = 0*.*29*, sign test). Cumulative density function of log ratios in A-B combined across layers. Monocular and Binocular CDFs significantly different (*p*<*0*.*001*, two-sided, two-sample Kolmogorov-Smirnov test). All recordings from awake mice trained in visual detection tasks.

### Spikes in binocular V1 show more OFF dominance than in monocular V1

Action potentials evoked during these same recordings showed weaker but consistent binocular OFF dominance and monocular ON dominance. We separated neurons into regular spiking (RS) and fast spiking (FS) units using peak-trough waveform width (Fig. S3; see Methods) and focused our analysis on putative excitatory RS neurons across all layers. In binocular V1, RS neurons showed significantly greater OFF dominance than those in monocular V1 (Fig. 3A; *p = 0*.*02*, two sided Kolmogorov-Smirnov test). Within each population, individual neurons in binocular V1 showed a mixture of OFF versus ON dominance (47% with log ratios < 0; n = 104; *p = 0*.*78*, sign test), while monocular RS neurons trended towards ON dominance (62% with log ratios > 0; n = 98; *p = 0*.*11*, sign test). We wondered if differences between the strength of ON and OFF dominance in spiking versus LFP resulted from the sparse and binary nature of awake spike responses (versus continuous LFP signals), so we turned to a larger data base of spike recordings next.

**Fig. 3.**
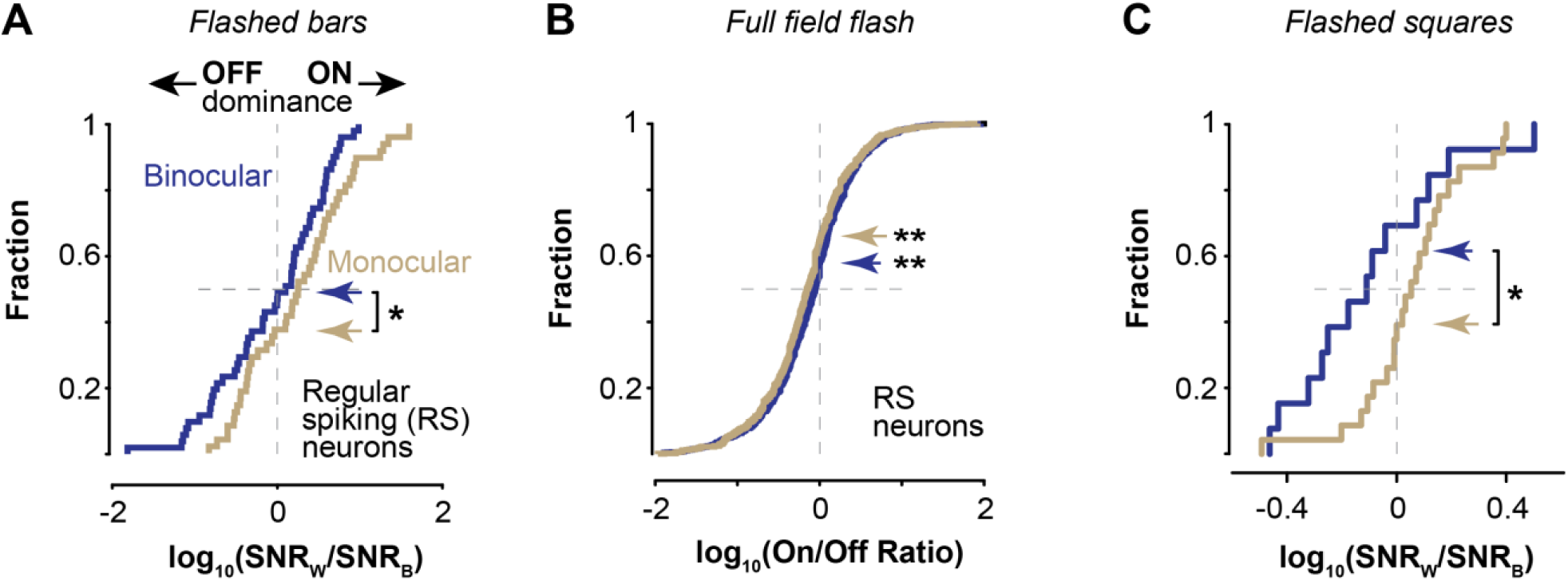
Spikes in binocular V1 show more OFF dominance than in monocular V1. **A:** Regular spiking neurons (RS) in binocular V1 (blue, n = 104) significantly more OFF dominant than in monocular V1 (gold, n = 98; *p = 0*.*022*, two-sided, two-sample Kolmogorov-Smirnov test). Monocular population trending towards ON dominance (log ratio: 0.26 ± 0.64; mean ± SD; 62% of data > 0; *p =0*.*11*). Binocular neurons not biased as a population (log ratio: −0.06 ± 0.65; mean ± SD; 47% of data < 0; *p =0*.*78*). Same experiments, mice, and stimuli as Figs. 1 – 2. See Fig. S4 for FS neurons. **B**. RS neurons in binocular V1 from Allen Brain Institute Neuropixels dataset significantly OFF dominated (n = 978; 54% of data < 0; *p* < *0*.*001*, sign test), as were monocular V1 neurons (gold, n = 834; 56% of data < 0; *p* < *0*.*001*, sign test). No significant difference between distributions (*p =0*.*78*, two-sided, two-sample Kolmogorov-Smirnov test). Responses evoked by full field black or white flash. Units grouped by azimuth followed by log_10_ transform of On/Off ratio (see Methods). See Fig. S4 for FS neurons. **C:** Responses to flashed squares (10°, 0.1s) in binocular V1 significantly more OFF dominated (n = 13; 69% of data < 0) than monocular V1 (n = 23; 61% of data > 0; *p = 0*.*031*, two-sided, two-sample Kolmogorov-Smirnov test). Full contrast black or white squares projected onto spherical half dome (see Methods) in awake mice.

Populations of RS neurons recorded with Neuropixels probes also revealed OFF dominance in binocular V1. We analyzed the publicly available Allen Brain Institute Visual Coding - Neuropixels database (Siegle et al., 2019). We first grouped neurons by azimuthal RF location (mapped with small grating patches), and then identified RS versus FS neurons according to spike waveform widths (see Methods). We separated RS neurons with RFs in the monocular versus binocular visual field, and then examined responses to full screen black or white flashes (0.25s duration, 1 s blank grey screen background). Binocular RS neurons (n = 978) exhibited clear preferences for black stimuli: the On/Off response ratio was significantly <1 (0.88 ± 1.6; median ± MAD; *p*<*0*.*001*; sign test; see Fig. S4D), and the log_10_ transformed ratios (just as calculated in Fig. 3A) were significantly OFF dominated (Fig. 3B; 54% of neurons with log ratios < 0; *p* < *0*.*001*, sign test). Unexpectedly, monocular RS neurons (n = 834) also exhibited significant OFF dominance to full screen flashes: the On/Off ratio was significantly less than 1 (0.81 ± 1.7; *p*<*0*.*001*; sign test; see Fig. S4), and the log_10_ transformed ratios were OFF dominated (56% of data < 0; *p* < *0*.*001*, sign test). However, there was no significant difference between the cumulative fractions of OFF dominance across the two groups (*p = 0*.*78*, two-sided, two-sample Kolmogorov-Smirnov test). FS neurons showed mixed trends (Fig. S4). Importantly, visual responses in this dataset were driven by sustained, full field black or white flashes, whereas our earlier results were obtained with brief, spatially localized bars (Fig.1-2, 3A). This difference suggests that ON versus OFF dominance in mouse V1 may also depend upon stimulus features (Yeh et al., 2009; Jansen et al., 2019), so we next examined an independent data set that also used sparse, spatially localized stimuli.

Spikes evoked by small, brief black or white squares also showed binocular OFF versus monocular ON dominance. In an independent set of experiments that projected visual stimuli (10° black or white squares, 0.1 s duration) onto a demispherical dome (Lopes et al., 2020), we analyzed visual responses in single units that showed high SNR and reliable receptive field maps (see Methods); this allowed us to group units according to azimuthal RF location. Consistent with results using briefly flashed bars (Fig. 3A), briefly flashed squares evoked significantly more OFF dominated responses in binocular versus monocular V1 (Fig. 3C; n = 36; *p = 0*.*031*, two-sided, two-sample Kolmogorov-Smirnov test). These complementary results (Fig. 3A, C) were obtained with similar recording techniques and stimulus conditions. Taken together, these three independent datasets of spiking activity reveal that binocular V1 consistently shows strong OFF dominance, while monocular V1 shows ON dominance when driven by brief, spatially localized visual stimuli, consistent with prior work (Polack and Contreras, 2012). We also examined population responses in simultaneously recorded RS ensembles (combining all RS neurons into single population RFs), and these revealed similar if not stronger trends, particularly towards monocular ON dominance (Peak white SNR = 9.7, black = 6.3; not shown). We next investigated if subthreshold synaptic inputs of binocular versus monocular RS neurons could explain OFF versus ON dominance.

### Spatially distinct OFF versus ON dominance in membrane potential of V1

Subthreshold membrane potential (V_m_) showed clear OFF dominance for binocular neurons, and ON dominance in monocular neurons. Using the exact same black or white bar stimuli that drove LFP (Figs. 1-2) and spikes (Fig. 3A), we measured V_m_ and spikes with whole-cell patch-clamp recordings in L2/3 (both awake and anesthetized recordings; see Methods). Neurons with RFs in binocular V1 (n = 13) showed depolarization to bars appearing in the binocular visual field (Fig. 4A, B), but responses to black bars evoked greater depolarization and significantly greater peak SNR (4.2 ± 0.1) than responses to white bars (3.4 ± 0.1, *p* < *0*.*01*, Wilcoxon rank sum test; Fig. 4C). In contrast, neurons with RFs in monocular V1 (n = 19) showed much larger depolarization to white versus black bars appearing in monocular visual space (Fig. 4D, E), and with a significantly larger SNR for white (3.3 ± 0.03) versus black stimuli (2.2 ± 0.02; *p* < *0*.*001*, Wilcoxon rank sum test; Fig. 4F). The log SNR ratios for V_m_ responses revealed clear OFF dominance for binocular neurons (77% of neurons with ratios < 0; n = 13), and ON dominance for monocular neurons (68% with ratios > 0, n = 19; *p* < *0*.*05*, two sample Kolmogorov-Smirnov test). These trends in V_m_ were even more pronounced for neurons that emitted spikes to at least 1 of the stimuli presented in the center ± 1 locations of the RF (Fig. 4H; 100% of binocular neuron V_m_ SNR ratios OFF dominated; 66% of monocular neuron V_m_ SNR ratios ON dominated; *p* < *0*.*001*, two sample Kolmogorov-Smirnov test). Finally, the spike responses in the same neurons (Fig. 4I; see also Fig. S5) revealed clear separation of OFF dominant responses in binocular V1 neurons, and ON dominated responses in monocular V1 neurons, both driven by the underlying selectivity of synaptic activity (Fig. 4I; *p* < *0*.*05*, two sample Kolmogorov-Smirnov test).

**Figure 4.**
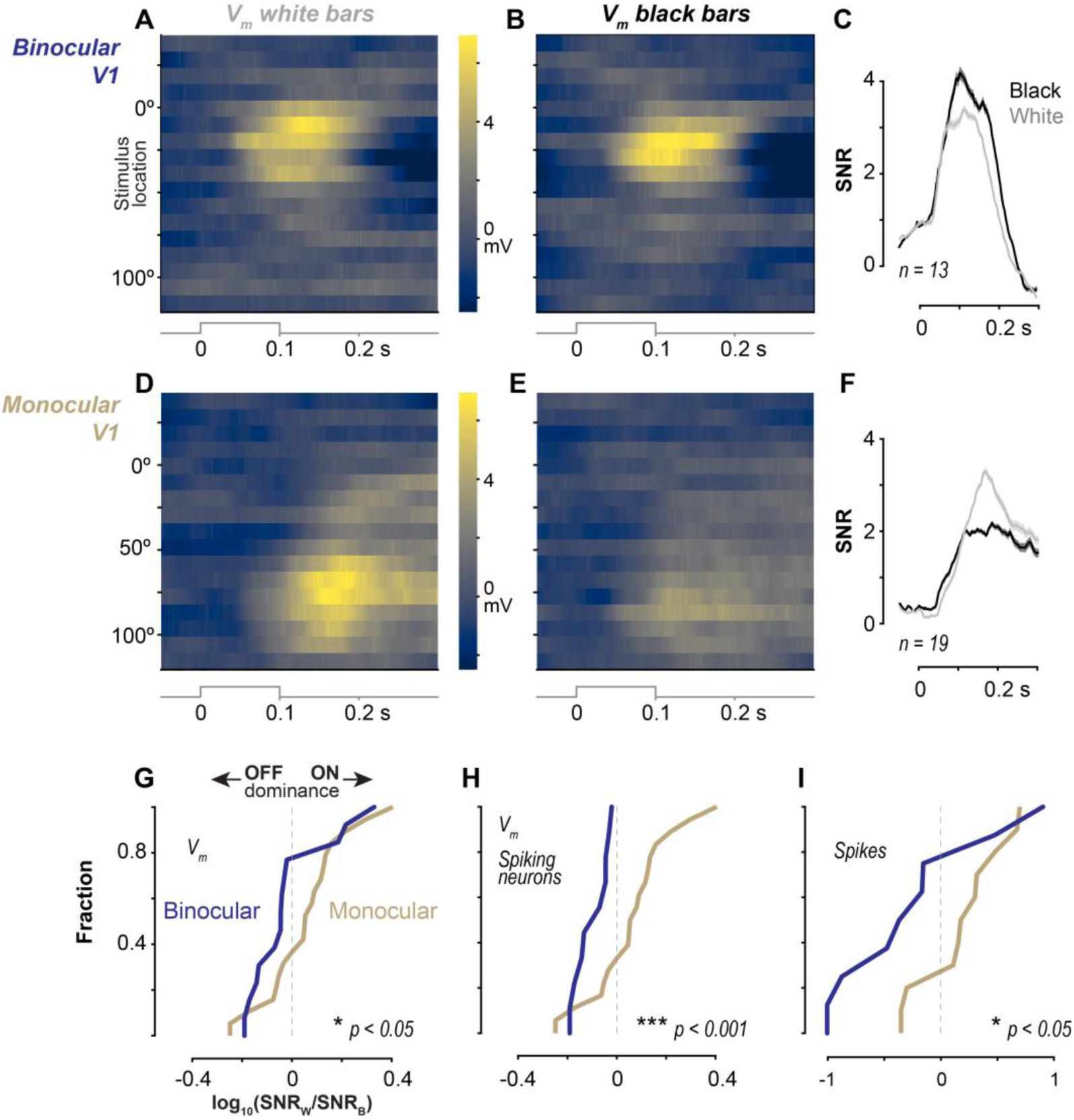
Membrane potential OFF dominance in binocular V1, ON dominance in monocular V1. **A:** Binocular V1 L2/3 membrane potential (V_m_) responses to white bars (n = 13). Peak V_m_ depolarization (Δ6.3 mV) centered at 10°. Average pre-stimulus V_m_ subtracted from each neuron before averaging. **B**. Same neurons as A, responses to black bars. Peak V_m_ depolarization (Δ6.7 mV) centered at 19°. **C:** Binocular V1 V_m_ signal to noise ratio (SNR, see Methods) for black versus white bars at center (± 1 locations) of receptive field (RF). Peak black SNR = 4.2 ± 0.1, white = 3.4 ± 0.1 (*p* < *0*.*01;* Wilcoxon rank rum; mean ± SEM; n = 13 neurons). **D-F**. As in A-C, for Monocular V1 V_m_ response (n = 19 neurons). Peak depolarization for white bars (Δ6.8 mV) larger than for black bars (Δ3.3 mV), both centered at 77°. Peak black SNR = 2.2 ± 0.1, white = 3.2 ± 0.1 (*p* < *0*.*001*; Wilcoxon rank rum). **G**. Log SNR ratios (see Methods) for binocular (blue) versus monocular (gold) neurons. Binocular neurons significantly more OFF dominated (77% with ratio <0) than monocular neurons (32% < 0; *p* < *0*.*05*, two sample Kolmogorov-Smirnov test). **H**. Same as G, for Vm log SNR ratio of spiking neurons (n = 9 Binocular, n = 18 monocular). Binocular neurons significantly more OFF dominated (100% with negative ratio) than monocular neurons (33%; *p* < *0*.*001*, two sample Kolmogorov-Smirnov test). **I**. Same as G, for spikes. Binocular spike responses significantly more OFF dominated (75% with negative ratio) than monocular neurons (20%; *p* < *0*.*05*, two sample Kolmogorov-Smirnov test).

## Discussion

Here we revealed that in mouse V1 the central visual field is strongly OFF dominated, while the peripheral visual field is more ON dominated. This spatial relationship pervaded multiple levels of neural activity in several independent data sets and was most prominent in LFP and membrane potential (V_m_). Since both LFP and V_m_ reflect population activity, this suggests that spatial OFF versus ON dominance in V1 emerges from large networks of neurons with shared luminance polarity preferences in specific retinotopic locations. Our findings help resolve prior conflicting results, and suggest common organizing principles for spatial processing of luminance increments and decrements in mammalian visual systems.

Our findings of OFF and ON dominance in mouse V1 echo and consolidate much prior work. Pioneering studies in cat and primate V1 showed OFF dominance for stimuli in the central portions of the binocular visual field (Jin et al., 2008; Yeh et al., 2009; Xing et al., 2010; Jin et al., 2011). Moreover, thalamic input to binocular V1 in cats is strongly OFF dominant, but ON contributions increase for peripheral stimulus locations (Jin et al., 2008). In ferret V1, stimuli presented peripherally elicit ON dominance (Smith et al., 2015). Likewise, wide-field voltage dye responses in mouse V1 show ON dominance for stimuli appearing in the monocular visual field (Polack and Contreras, 2012); however, Ca^2+^ responses from mouse V1 neurons with RFs in binocular V1 (20–35° azimuth) show OFF dominance (Jimenez et al., 2018). Accordingly, neurons spanning binocular and monocular regions of mouse V1 show equal ON and OFF dominance as measured with Ca^2+^ responses (Smith and Hausser, 2010). Our results with electrophysiological recordings across multiple levels not only consolidate and explain these discrepancies in mice but reveal broad consistency with observations in other species. Together, our results suggest a potentially simple organizing principle for mammalian visual systems: OFF and ON dominance coexist but vary with retinotopic position.

In response to brief spatially localized stimuli, LFP, spikes, and V_m_ all showed strong binocular OFF dominance, but generally weaker monocular ON dominance. Although monocular OFF dominance was unexpectedly observed in the Allen institute Neuropixels data set, this could perhaps be explained by the stimulus: a sustained full-field flash. Indeed, as stimuli become larger or lower spatial frequency, OFF dominance increases (Jansen et al., 2019). Future experiments could test this stimulus dependence, with a prediction that monocular responses gain OFF dominance as stimulus size increases or spatial frequency decreases.

We found that OFF or ON dominance permeated all cortical layers at a given spatial location. This is different from primates, where strong OFF dominance emerges in L2/3. This may be related to exquisite laminar organization of thalamic inputs to V1 in primates (Callaway, 1998) versus less delimited projections in mice (Antonini et al., 1999). Further, mouse V1 lacks the organizing structure of maps of ocular dominance, orientation, and ON/OFF subfields, as seen in cats, primates, and other mammals (Smith et al., 2015; Lee et al., 2016). Since we observed strong binocular OFF dominance and weaker monocular ON dominance in L2/3 as well as L4, this suggests thalamic projections may seed binocular versus monocular V1 with OFF versus ON biases; indeed, these spatial biases may arise in mice from spatial gradients of ON versus OFF retinal ganglion cells (Bleckert et al., 2014; Schroder et al., 2020). Importantly, despite the lack of orientation maps in mouse V1, retinotopic maps are clear; moreover, L2/3 neurons <200 microns from one another share highly localized and overlapping ON and OFF subfields (Smith and Hausser, 2010), providing a substrate for coherent local population responses to bright versus dark stimuli that were visible in the LFP and V_m_ of L2/3.

We revealed that subthreshold synaptic responses in mouse V1 are selective for luminance polarity as a function of retinotopy. Subthreshold selectivity was most pronounced for cells that spiked. ON versus OFF selectivity may be amplified by spike threshold, as with many other visual computations (Priebe and Ferster, 2012). Importantly, patch-clamp recordings report every single spike in sparse firing conditions, and allow direct comparison to the selectivity of V_m_. Furthermore, we measured clear and strong spatial dependence for ON versus OFF dominance in both Vm and LFP, in entirely separate mice and experiments. Given the close relationship between LFP, V_m_, and synaptic activity in mouse V1 (Haider et al., 2016), this suggests that presynaptic populations share coherent selectivity for both stimulus position and luminance polarity preference. This could ensure that ON and OFF computations in V1 also contain appropriate spatial signals for downstream targets.

ON versus OFF dominance was less pronounced in extracellular spikes than LFP or V_m_, for several potential reasons. First, black or white bars appeared at a single (vertical) orientation that was likely sub-optimal for driving spikes in most neurons. However, even sub-optimal stimuli depolarize membrane potential (Priebe and Ferster, 2012; Lien and Scanziani, 2013), and LFP responses summate subthreshold activation across local neural populations (Katzner et al., 2009). This may explain why ON and OFF dominance trends were most visible for these signals. Despite this limitation, four independent sets of spike measurements (3 extracellular; 1 intracellular) showed spatially separated OFF versus ON dominance of spiking, consistent with the trends in LFP and V_m_. Although monocular and binocular extracellular spike distributions were often not themselves significantly ON or OFF dominant, the two distributions were largely separated from one another. Lastly, unlike the inherently correlated population activity underlying ON or OFF dominance in LFP and V_m_ responses, metrics calculated from spikes did not consider local population correlations, a topic for further study.

Finally, why might binocular OFF versus monocular ON dominance be beneficial for mice? During navigation, rodents prioritize binocular visual field coverage (Wallace et al., 2013; Meyer et al., 2020), perhaps underlying computations of self-motion versus visual motion (Saleem, 2020). Drifting versus looming dark stimuli in binocular visual space elicit fleeing versus freezing (Wallace et al., 2013; De Franceschi et al., 2016). Additionally, mice use binocular vision to perceive depth (Samonds et al., 2019), to hunt for prey (Hoy et al., 2016; Shang et al., 2019; Michaiel et al., 2020), and forage during daylight (Daan et al., 2011; Hut et al., 2011). While stationary, mice show greatest perceptual sensitivity for stimuli in binocular visual space (Speed et al., 2019), but can use spatial attention to improve perceptual sensitivity for stimuli in monocular visual space (Speed et al., 2020). Thus, mice show several specializations in spatial visual processing and behavior commonly found in other mammals, including spatially specific OFF versus ON processing as shown here. Techniques in mice will enable detailed investigation of the circuits and mechanisms driving a variety of spatial visual behaviors elicited by luminance increments and decrements, a fundamental feature of vision.

## Methods

All procedures were approved by the Institutional Animal Care and Use Committee at the Georgia Institute of Technology and University College London and were in agreement with guidelines established by the National Institutes of Health, and The Animal (Scientific Procedures) Act 1986.

### Experimental model and subjects

Recordings in Haider lab used male and female mice. Recordings from the Allen Institute – Neural Coding database are fully detailed elsewhere (Allen Brain Observatory, 2019). Both male and female mice were used, and only one mouse was used per recording. Recordings in Saleem lab used male mice.

*See Data Tables 1 – 3 at end of Methods for full details on genotypes, mouse numbers, and experimental sampling for each set of recordings*.

**Table 1.** Experimental subjects. Haider lab

**Tables 1 – 3. Experimental subjects**
| *Table 1. Haider lab*<br>**Mouse Strain** | **n = mice [silicon probes; patch clamp]** | **n = recordings [silicon probes; patch clamp]** | **RRID** |
| --- | --- | --- | --- |
| B6PV<sup>Cre</sup> | [3;1] | [31;1] | IMSR_JAX:017320 |
| C57BL/6J | [3;0] | [19;0] | IMSR_JAX:000664 |
| Ai32 | [1;0] | [13;0] | IMSR_JAX:024109 |
| Sst-IRES-Cre | [1;1] | [5;2] | IMSR_JAX:013044 |
| Ai32 x B6PV<sup>Cre</sup> | [3;7] | [26;10] | See above |
| Ai32 x Sst-IRES-Cre | [3;2] | [62;8] | See above |
| Ai32 x Scnn1a-cre | [0;4] | [0;7] | IMSR_JAX:009613 |
| CNTNAP2<sup>-/-</sup> | [0;1] | [0;1] | IMSR_JAX:017482 |
| Ai40 | [0;1] | [0;1] | IMSR_JAX: 021188 |
| Uncertain genotype | [0;1] | [0;2] | -- |

**Table 2.** Experimental subjects. Saleem Lab

| *Table 2. Saleem Lab*<br>**Mouse Strain** | **n = mice** | **n = recording sessions** | **RRID** |
| --- | --- | --- | --- |
| C57BL/6J | 6 | 25 | See above |

**Table 3.** Experimental subjects. Allen Institute

| *Table 3. Allen Institute*<br>**Mouse Strain** | **n = mice** | **n = neurons** | **RRID** |
| --- | --- | --- | --- |
| C57BL/6J | 26 | 1444 | See above |
| Ai32 x B6PV<sup>Cre</sup> | 5 | 195 | See above |
| Ai32 x Sst-IRES-Cre | 10 | 504 | See above |
| Ai32 x Vip-IRES-Cre | 6 | 347 | See above |

### Surgical preparation and neural recordings

#### Haider lab

Detailed methods have been described previously (Speed et al., 2019). Mice (5 – 8 weeks old; reverse light cycle individual housing; bred in house) were chronically implanted with a stainless-steel headplate with a recording chamber during isoflurane (1-2%) anesthesia. After implant surgery mice recovered for 3 days before experimentation. All silicon probe recordings here were from mice that were trained over several weeks to perform a visual spatial detection task (Speed et al., 2020). At the conclusion of training, a small craniotomy (100-400 microns) was opened over monocular or binocular V1 during isoflurane anesthesia. Mice were allowed ≥3 hours of recovery before awake acute recordings. Single shank linear 32 site silicon probes (Neuronexus, A1×32) were used to record neural activity across cortical layers. The electrode was typically advanced to 1000 microns below the dura, and the site was covered in sterile artificial cerebrospinal fluid (aCSF). After the electrode settled (∼15 minutes), mice performed a visual spatial detection task for ∼ 2 hours (Speed et al., 2020). At the end of the behavioral sessions, mice were presented black and white visual stimuli to map spatial receptive fields (described below); we focused on these data sets recorded at the end of behavioral sessions in trained mice to limit the effects of spontaneous behavioral variability in mice not performing visual tasks. Nevertheless, major findings regarding ON and OFF dominance from mice trained in behavioral tasks (Fig. 1, 2, 3a) were also evident in untrained mice (Fig. 3b-c; 4). As a control for any possible effects of training, combining recordings from both trained and passive awake mice (n = 94) did not diminish strong OFF dominance in binocular V1 (*p* < *0*.*001;* sign test), and the trend towards ON dominance in monocular V1 was even stronger (n = 50; *p = 0*.*04*, sign test). Whole-cell patch-clamp recordings were performed in awake (n = 9) and anesthetized mice (n = 10), as detailed previously (Haider et al., 2013). We observed no significant differences in ON/OFF dominance within spatial location for awake versus anesthetized V_m_ recordings, so these were combined within location (awake vs. anesthetized SNR log ratios monocular: 0.07 ± 0.1 vs 0.05 ± 0.2, *p = 0*.*6*; binocular: −0.04 ± 0.2 vs −0.05 ± 0.2; *p = 0*.*8*, Wilcoxon rank sum tests; median ± IQR).

#### Saleem lab

Detailed methods have been described previously (Lopes et al., 2020). Mice were implanted with a custom-built stainless-steel metal plate under isoflurane anesthesia. The area above the left visual cortex was kept accessible for electrophysiological recordings. Seven days following the surgery mice underwent the first habituation session in the virtual reality apparatus. Following the habitation period (one session per day, 8-13 days), a craniotomy over V1, centred at (2 mm lateral to sagittal midline and 0.5 mm anterior to lambda) was performed. Mice recovered for 4-24 hours before the first electrophysiology recording session. Multiple recording sessions were executed from each animal (one per day, n = 37 recordings, min 2, max 9). To preserve the brain tissue we left the dura intact. This was pierced locally by the silicon probe (ASSY-37 E-1, Cambridge Neurotech Ltd.) at the beginning of each recording session. Mice were free to run while presented black and white visual stimuli to map spatial receptive fields (described below).

### Visual stimuli

#### Haider lab

Mice were shown vertical bars at various contrasts and spatial locations (Fig. 1). Bars were 9° wide and covered the whole height of the screen (spanning 50°). Bars were shown at 17 locations covering binocular and monocular areas of visual space, spaced evenly every 9.6 ° from −37.8° to 115.8° azimuth. The vertical meridian was defined as 0°. Contrasts ranged from 100% black to 100% white. Pixel values ranged from 0-255, and grey was set at 128. Michaleson contrast was calculated as percent pixel value difference from grey background with the following equation

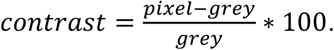

Each stimulus lasted 0.1s before disappearing, and subsequent stimuli appeared after 0.3s of grey screen. Stimuli were shown 10 times for each polarity, contrast, and location. Contrasts levels were 5, 10, 25, 40, 50, 75, and 100% for both black and white, but not all contrasts were shown in all sessions. Stimulus sequences were randomized for location, polarity, and contrast.

#### Saleem lab

Briefly, awake mice were shown a series of sparse noise frames consisting of a 9x#x00D7;9 (10° squares) or 8×8 grid (15° squares). The squares could each independently be white (pixel = 255), black (pixel = 0) or grey (pixel = 128). Only five squares could either be white or black at any frame presentation. Frames were shown for 0.1 s each in immediate succession. A single session lasted five minutes, or 3000 frames. Frame sequence was used posteriorly to construct a neural response to black, white, and grey stimuli in each square location.

#### Allen institute

Detailed stimulus parameters are described elsewhere (Allen Brain Observatory, 2019). Briefly, awake mice were shown a variety of stimuli in blocks. A receptive field mapping stimulus (20º drifting grating, 0.25s duration, 0.04 cycles per degree; 2 Hz temporal frequency) appeared at a single randomly chosen location on the screen (forming a 9 x 9 grid) tiling the whole visual field. Stimuli also included full-field flashes of black and white (0.25 s duration, 2 s inter-trial interval with uniform grey screen).

### Electrophysiological recording and analysis

#### LFP analysis (Haider lab)

Electrical signals were acquired through a Cereplex Direct (Blackrock Microsystems). Raw neural signals were acquired at 30 kHz. Local field potentials (LFP) were band pass filtered at 0.3-200Hz. Layers were identified via current source density analysis (Niell and Stryker, 2008; Speed et al., 2019, Figure S1).

#### Spike sorting (Haider lab)

Single unit activity was isolated with a semi-automated sorting algorithm (Rossant et al., 2016), as detailed in our previous studies (Speed et al., 2019). We classified single units as fast-spiking (FS, waveform peak-to-trough < 0.57ms) and regular spiking (RS, peak-to-trough > 0.57 ms) based on their waveform widths (Figure S3). FS neurons in mice are predominantly paravalbumin (PV) positive inhibitory neurons, while >85% of RS neurons are putative excitatory neurons (Speed et al., 2019).

#### Vm analysis (Haider lab)

Detailed methods have been described previously (Haider et al., 2013; Speed et al., 2020). Whole-cell patch-clamp recordings were performed in current clamp mode (Molecular Devices, Multiclamp 700B) and acquired at 20 kHz with custom software (MATLAB). All recordings were in L2/3 based on depth estimated from the micromanipulator. Visual stimuli were displayed as described above, but only at 100% contrast. In some cases neurons only emitted spikes for bars of one color, so calculations for log ratios were assigned SNR = 1 for the bar color with no spikes.

#### Spike sorting (Saleem lab)

Single units were isolated using a semi-automated algorithm (Pachitariu et al., 2016). Then, we selected only units that had a mean firing rate greater than 0.5 Hz in the first and last third of the sparse noise presentation period.

#### Spike sorting (Allen Institute)

Neurons in the Allen Institute dataset were pre-sorted and packaged with several pre-computed quality metrics, as detailed elsewhere (Allen Brain Observatory, 2019). We plotted histograms of spike waveform widths, and observed a clear bimodal distribution with a natural partition at 0.42 ms, and used this to classify RS and FS groups.

### Receptive field (RF) map analysis

#### Haider lab

Recording sites in V1 were targeted with stereotaxic coordinates and/or intrinsic signal imaging, and further verified with functional localization of visual spatial receptive fields. A receptive field (RF) map for each recording session was created by first averaging together the LFP responses per electrode channel across all stimulus contrasts per stimulus location. This resulted in a 3D matrix of LFP responses: [stimulus location x time x probe channel]. The median response across probe channels (averaging across laminar depth) generated a global map of spatial responses for each azimuth location across time. The stimulus position that evoked the largest LFP activation (depth negative voltage response; see Fig. 1) was designated as the central location of the receptive field (RF). Recordings with central RF locations < 40° in azimuth were classified as binocular recordings (median RF preference: 19.8 ± 7.1°, n = 103) whereas those with preferred location > 55° were classified as monocular (median RF preference: 77.4 ± 8.5°, n = 50). This categorization is broadly consistent with anatomical and physiological definitions of the binocular and monocular representations in mouse V1 (Drager, 1975; Niell, 2015). Here we focused on responses elicited by 100% contrast black or white stimuli, since these were the most reliable and comparable across recordings (Fig. S2). For spiking data, RF maps were calculated as described above per neuron. Singular Value Decomposition (SVD) was used to automate the RF localization process. SVD reduce RF maps to two 1-D components, and the maximum value on the spatial component was designated as the central RF location. These single neuron RF locations were verified by comparison to the RF location reported by the LFP during the same recordings. In some analyses, population RF maps were created by combining single unit RF maps recorded simultaneously per recording (within visual area and cell type).

#### Saleem lab

Receptive field mapping was performed in awake mice free to run or rest on a treadmill (polystyrene wheel) in a virtual reality environment (Lopes et al., 2020). Black and White squares were projected in a 2-D grid pattern Inside a hemispherical dome that spanned 240° in azimuth and 120° in elevation on the right visual field. For a particular square stimulus location and color (i.e., a frame), a sliding window of 100 ms was used to bin spikes into discrete time points relative to stimulus onset. Time bins started at 10 ms prior to stim onset, and ended at 120 ms after stimulus onset, in intervals of 10 ms (14 time points). For example, the bin starting at 50ms would contain all spikes from 50-150 ms relative to stimulus onset. At each time point, an RF map (see above) was created by combining the responses to all squares in the 9×9 or 8×8 grid into one large response. For every neuron, the response map with the highest variance was selected in time – one for each black response and white response. These two maps were used for subsequent analyses. SVD was performed on each map, black and white independently, to determine the location of the best response in azimuth and elevation. Selection criteria were instituted specifically on this data set with 2-D stimuli. Neurons with firing rate <1 spike/s at the best location for both the black *and* white response were not analyzed. We next calculated for each neuron and for each stimulus color the percentage of locations in the RF map where normalized activation was >0.7 of the global max firing rate. Neurons where >40% of the map exceeded this threshold in both black and white response (i.e., there was no spatially localized RF) were excluded. Visual inspection of included neurons showed clear RFs. Stimulus locations where normalized activation was >0.9 of the global max response were all expected to be within 1 location (+/-10°) of the maximum response location. Neurons with normalized activation >0.9 of the global max response at non-contiguous spatial positions in *both* black and white RF maps were excluded. We note that conditioning selection criteria for the combined ON and OFF responses prevented exclusion of neurons that were strongly responsive to one color, but not at all to the other color. We then calculated SNR and log_10_ transformed ratios as described previously, sorting neurons into binocular versus monocular groups by best azimuthal RF locations, using the same criteria as before.

#### Allen Institute

This data set was pre-processed by the Allen Institute to pass several quality metrics and quantify visual feature selectivity. The data set precomputes optimal azimuthal receptive field location for each neuron, which we used to separate cells into binocular (−45 to 45°) and monocular (55° to 130°) groups. Full-field black or white flashes were used to calculate the ON/OFF response ratio for each cell as the mean firing rate during “ON” presentations divided by the mean firing rate during “OFF” presentations. A log_10_ transform was applied to these precomputed ratios just like all other data sets (see Fig. S4).

### Laminar identification

#### Haider lab

Laminar LFP responses were separated by using current source density analysis (Fig. S1), We defined L4 to span ± 100 microns around the location of the earliest and largest CSD sink, consistent with prior functional and anatomical localization of L4 in mouse V1 (Lien and Scanziani, 2013; Speed et al., 2020). After designating L4, the other layers (L2/3, L5/6) were analyzed by taking the median across channels within layer. Neurons were assigned to specific layers based on the location of the channel with the largest amplitude spike waveform. Spike analysis for all datasets combined neurons across layers because of unequal sampling, and because there was no clear indication of laminar differences in LFP analysis (Fig. 2).

### SNR calculation

We quantified ON versus OFF dominance using previously established methods (Yeh et al., 2009). Raw LFP and spiking traces were converted into signal-to-noise ratio (SNR) traces. SNR was calculated as

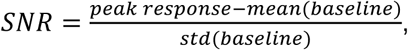

where baseline represents the response during the pre-stimulus period (global activity level in the −0.1s preceding stimulus onset for all stimulus locations). For single-unit data, if the standard deviation of the baseline equalled 0 (no activity), the SD (baseline) component was artificially set at 1. The peak response was generally restricted to a window spanning the earliest visual response latency (LFP data: 30-100ms after stimulus onset; spiking data: 0-180ms).The raw response was calculated differently for LFP vs. single-unit data. For LFP, the raw response was the mean of the preferred center location ±1 adjacent locations. Since spike RFs are narrower than LFP RFs (Haider et al., 2013), the peak response for spike RFs was calculated as the max (not mean) across the center ±1 locations. In all cases, conversion to SNR resulted in a trace of response amplitude normalized by the SD. To quantify responses to black and white stimuli, we computed a metric based on the log_10_ ratio of the responses to white stimuli versus black stimuli, consistent with previous studies (Yeh et al., 2009)

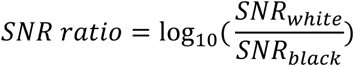

For each stimulus polarity, we found the max SNR value for both white and black stimuli, and computed SNR at this same latency for both stimulus polarities. Neurons in which either the white or black SNR was 0 were excluded (single polarity responders). The results were unaffected if we took the SNR max of each stimulus polarity at their respective peak latencies. SNR ratios < 0 were classified as OFF dominant, while SNR ratios > 0 were classified as ON dominant. For the Allen Institute data set, ON-OFF ratios were already pre-computed (Allen Brain Observatory, 2019), and these were used for the above equation.

## Funding

This work was supported by the Goizueta Foundation Fellowship (to J.D.R.), Alfred P. Sloan Foundation’s Minority Ph.D. (MPHD) Program Fellowship (to J.D.R.), the Whitehall Foundation (to B.H.), the Alfred P. Sloan Foundation Fellowship In Neuroscience (to B.H.), National Institutes of Health Neurological Disorders and Stroke (NS107968 to B.H.), National Institutes of Health BRAIN Initiative (NS109978 to B.H.), the Simons Foundation Autism Research Initiative (to B.H.), Sir Henry Dale Fellowship from the Wellcome Trust and Royal Society (200501 to A.B.S.), Human Science Frontiers Program grant (RGY0076/2018 to A.B.S).

## Data Availability

All data structures and code that generated each main figure will be deposited on Figshare.

## Author Contributions

B.W. built the data base, wrote analysis code, and analyzed all experiments; J.D.R., A.S., T.M. performed silicon probe experiments; S.C., E. K. M. performed patch-clamp experiments, B.W., S.C., T.M., L. M-B., B.H. performed data analysis; B.W., A. B. S., and B.H. wrote the manuscript with feedback from all authors.

## Supplemental Material

**FIGURE S1.**
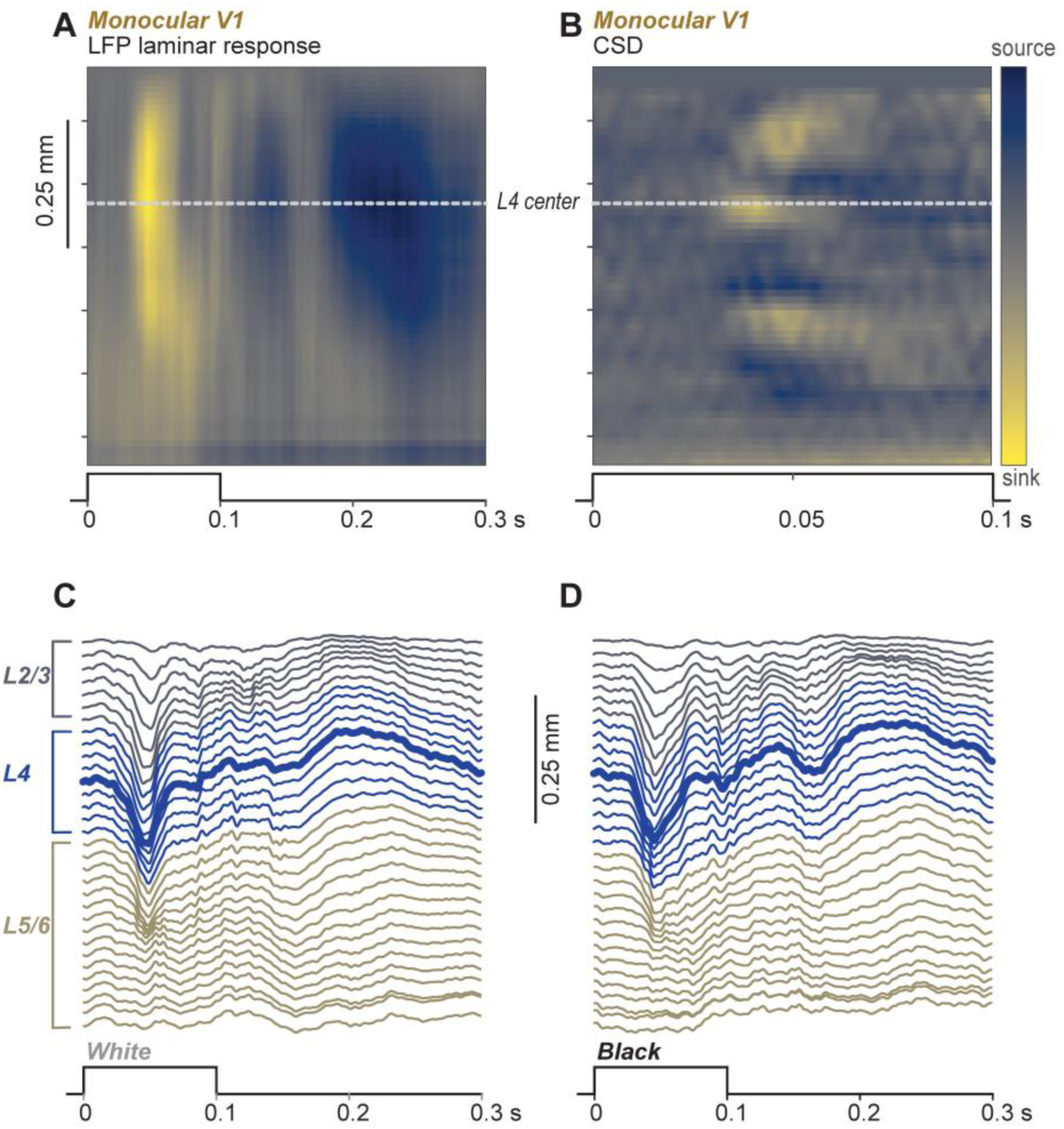
**A:** Average laminar LFP response in monocular V1 for black bars appearing at center of receptive field (RF). Same recording as **Fig. 1C**. Depth negative LFP (activation) in yellow. Stimulus timing shown below heat map. **B:** Current source density (CSD) map of laminar response in **A**. Center of Layer 4 (L4) identified by the earliest sink (dotted white line). Corresponding stimulus timing shown below heat map. Note that only 0.1 seconds is shown in CSD here for clarity, but CSD is calculated from 0-0.3 s post-stimulus. **C-D:** LFP traces from example recording in **A-B**, separated by layer. Stimulus timing shown below traces. Trial-average responses to 100% contrast white (**C**) and black (**D**) stimuli, at center of RF. Bold blue trace indicates center of L4. Response is stronger for white versus black stimuli.

**FIGURE S2.**
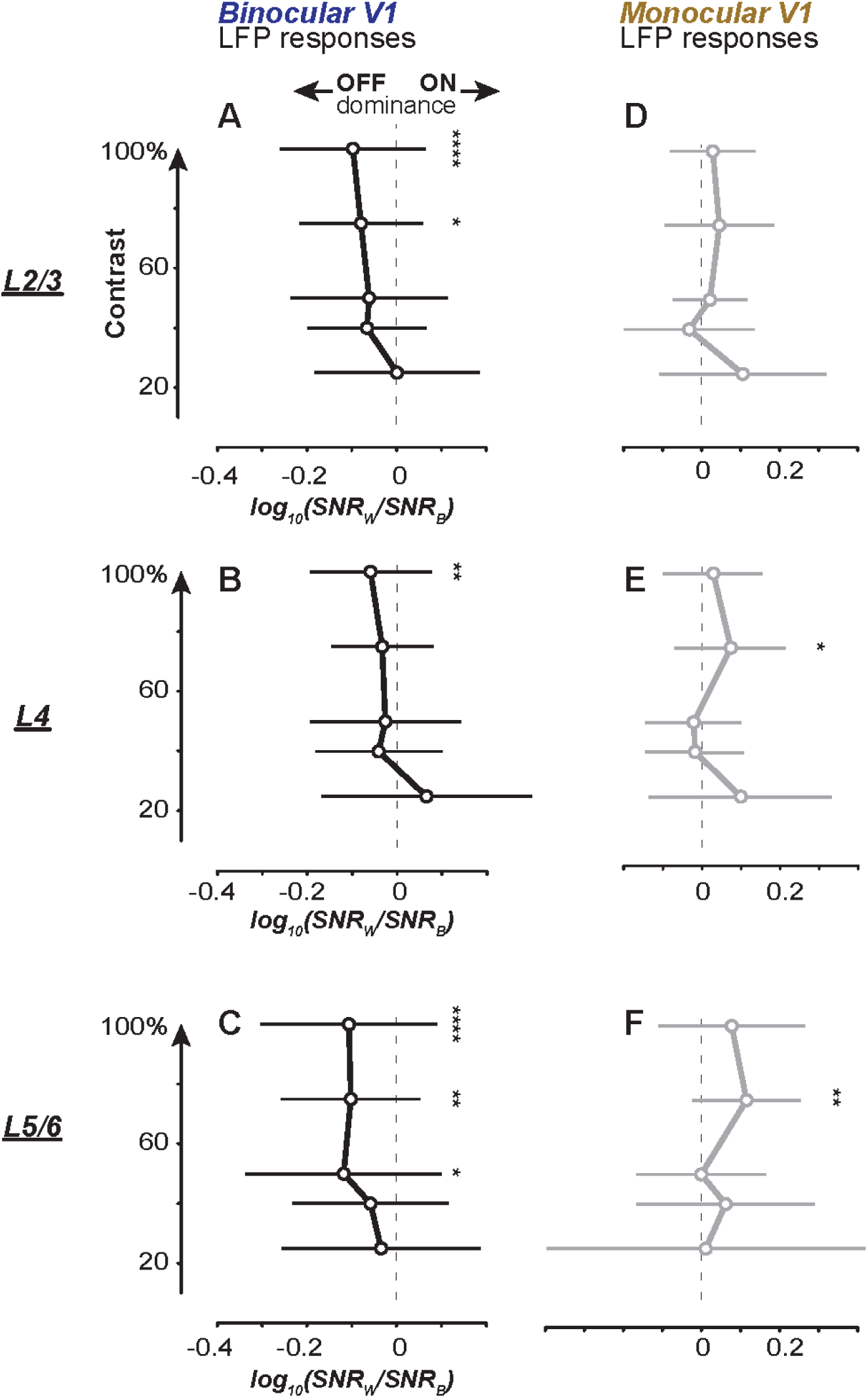
**A-C:** Log ratio of Binocular V1 LFP SNR for white vs. black stimuli of multiple contrasts. Error bars show mean ± SD response within layer 2/3 (L2/3, **A**), L4 (**B**), and L5/6 (**C**). Log ratio significantly negative at highest contrasts in all layers, and significant or trending negative across lower contrasts; negative value indicates OFF dominant response. P-values indicated by asterisks (*\* p*<*0*.*05; ** p*<*0*.*01; *** p*<*0*.*001; **** p*<*0*.*001*). **D-F:** Same as **A-C** for monocular recordings.

**FIGURE S3.**
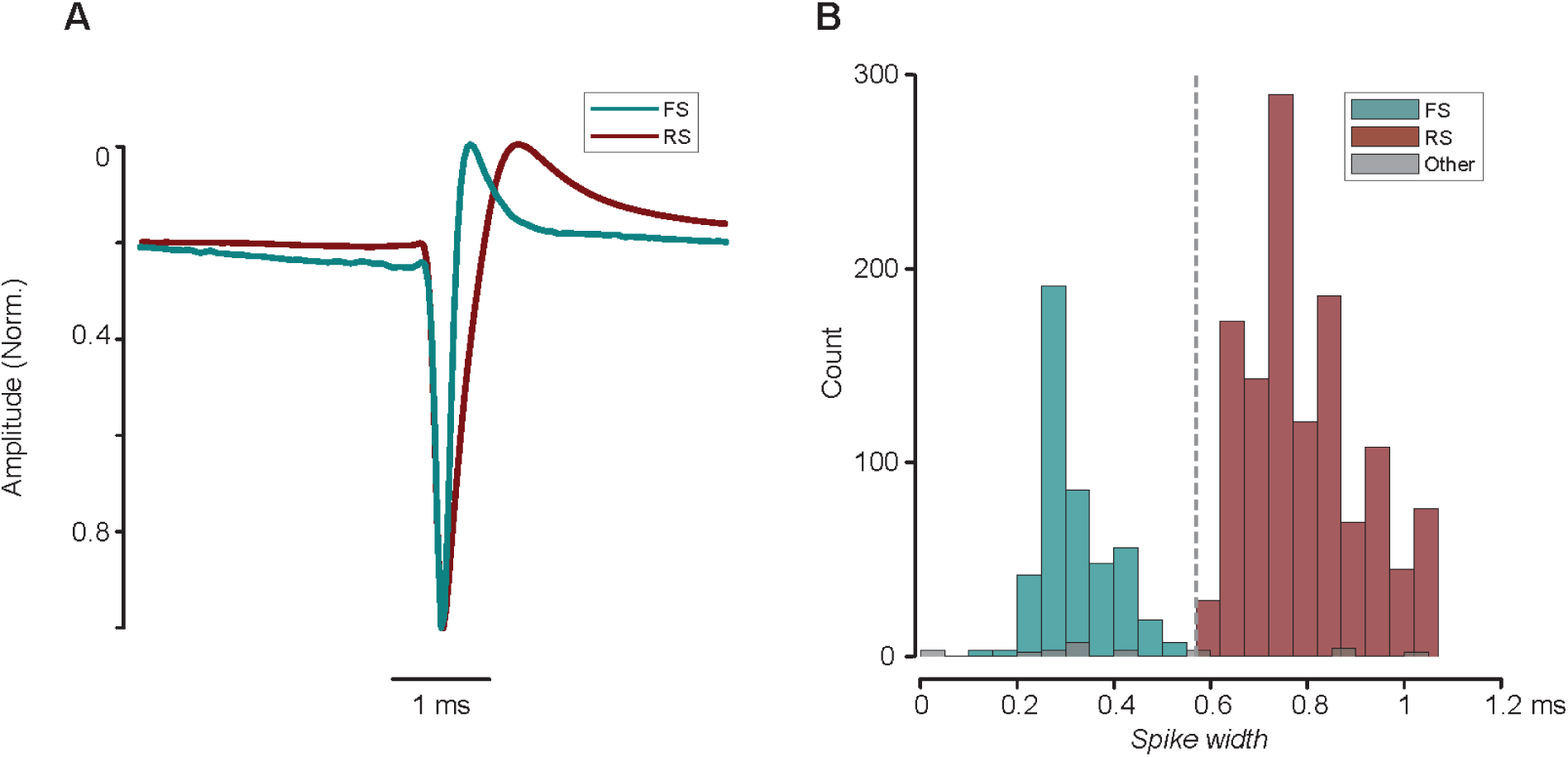
**A:** Average spike-triggered-average waveform for FS (turquoise) and RS (maroon) cells. (n=467, n=1240) shown in Fig. 3. **B:** Histogram of spike width distributions (full width at half height) for all FS (turquoise) and RS (maroon) cells. Note the clear separation at 0.57s (dashed line), consistent with prior studies (Speed et al., 2019; Niell and Stryker, 2010). Widths for cells that could not be robustly classified due to atypical waveform features in grey.

**FIGURE S4.**
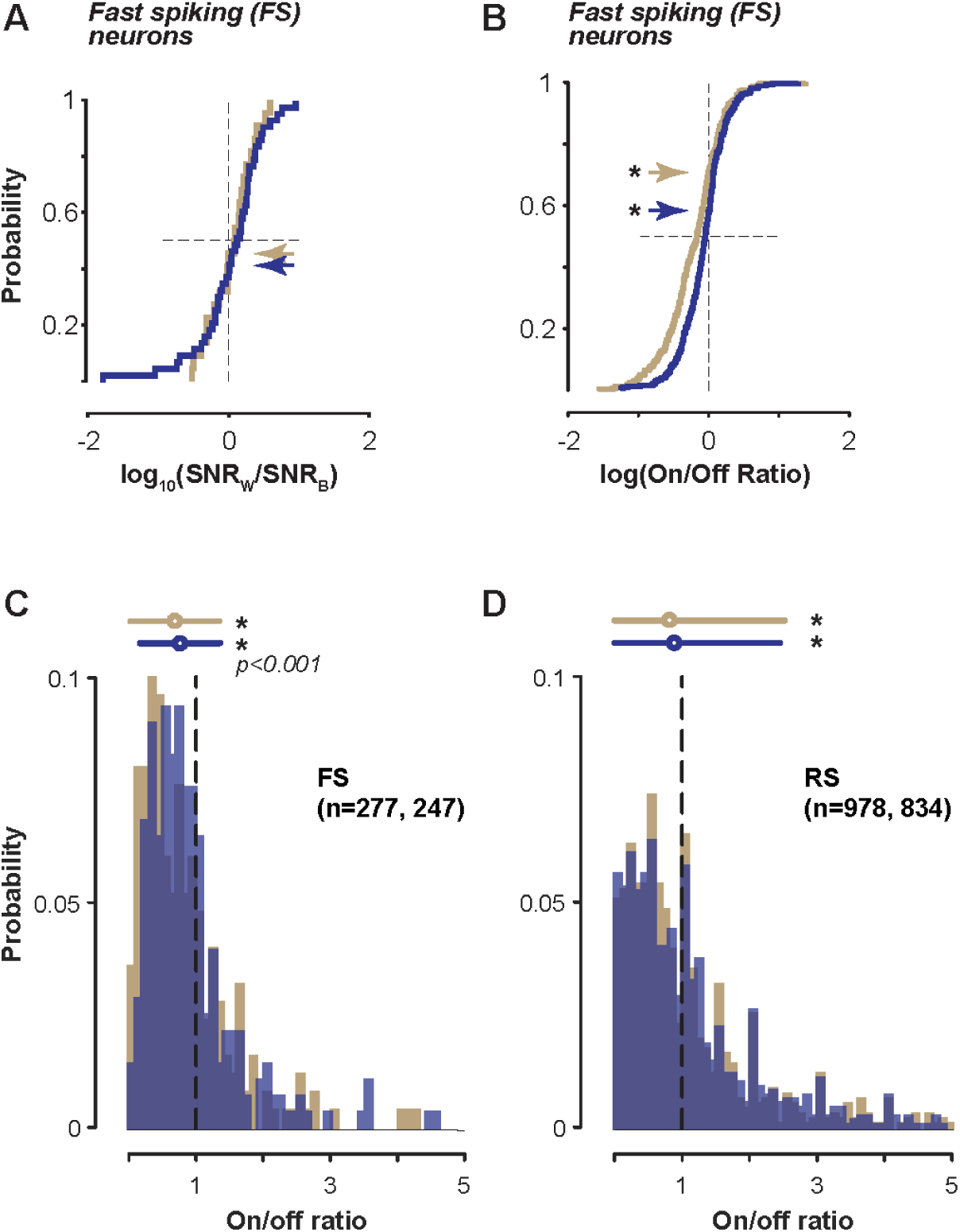
**A:** Silicon probe recordings of fast spiking (FS) neurons (same experiments as Fig. 3A). Binocular (blue) FS neurons (n = 88) split amongst ON and OFF dominant fractions (0.031 ± 0.49; mean ± SD; 40% of data < 0; *p = 0*.*22*). Monocular FS neurons (n = 46) also split between ON and OFF dominance (0.039 ± 0.32; 55% of data > 0; *p = 0*.*83*). **B:** FS neurons in Allen Brain Institute Visual Coding – Neuropixels dataset. 69% of binocular FS neurons and 70% of monocular FS neurons OFF dominated (*p* < *0*.*001* for both, sign test). Cumulative fraction of log transformed On/Off response ratios (see Methods). **C:** On/Off response ratios in FS neurons in Allen Brain Institute dataset. Binocular (blue, n = 277) and monocular (gold, n = 247) neurons ON/Off ratios significantly < 1 (Binocular: 0.76 ± 0.6; Monocular: 0.68 ± 0.66; median ± mad; *p*<*0*.*001* for both; sign test). Responses evoked by full field black or white flash (0.25s duration, 1 s interval). Units grouped by best azimuthal location to flashed grating patches (see Methods). Errorbars above distributions show median ± iqr. **D:** As in C, for Regular spiking (RS) neurons in binocular (blue, n = 978) and monocular (gold, n = 834) V1 significantly OFF dominated (Binocular ratio: 0.88 ± 1.6; Monocular ratio: 0.81 ± 1.7; *p*<*0*.*001* for both; sign test). Same data sets as C.

**FIGURE S5.**
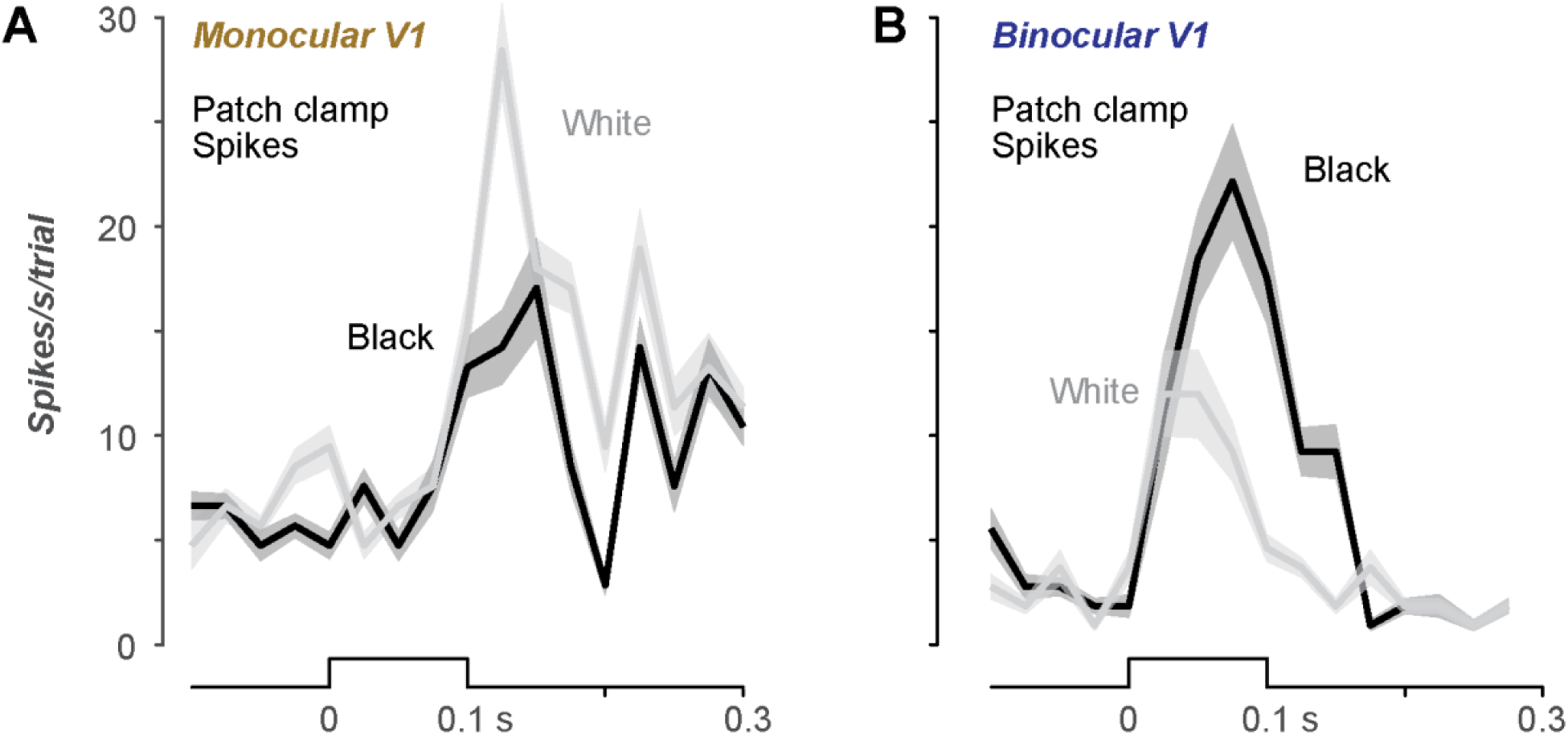
**A**. Spike responses in whole-cell patch-clamp recordings from L2/3 neurons in monocular V1 (Fig. 4). Spike density function across all cells (mean ±SEM) for responses to black or white bars in center±1 locations of RF. ON response is 67% larger than OFF response. **B**. Same as in A, for recordings in binocular V1. OFF response is 85% larger than ON response.

## Notes

### Competing Interest Statement

The authors have declared no competing interest.

